# Diverse responses among wild banana species to vapour pressure deficit, a solution for drought tolerance breeding?

**DOI:** 10.1101/2022.08.05.502925

**Authors:** David Eyland, Clara Gambart, Rony Swennen, Sebastien Carpentier

## Abstract

1

The predicted rise in global temperature is not only affecting plant functioning directly, but is also increasing air vapour pressure deficit (VPD). The yield of banana is heavily affected by water deficit but so far breeding programs have never addressed the issue. A reduction in transpiration at high VPD has been suggested as a key drought tolerance breeding trait to avoid excessive water loss, hydraulic failure and to increase water use efficiency. In this study, stomatal and transpiration responses under increasing VPD at the leaf and whole-plant level of 8 wild banana (sub)species were evaluated, displaying significant differences in stomatal reactivity. Three different groups were identified under increasing VPD. *M. acuminata* spp. e*rrans* (group I), *M. acuminata* spp. *zebrina* (group II) and *M. balbisiana* (group II) showed the highest transpiration rate limitations to increasing VPD. In contrast to group I, group II only showed strong reductions at high VPD levels, limiting the cost of reduced photosynthesis and strongly increasing their water use efficiency. Group II genotypes thus show favourable responses for high water use efficiency in regions with high VPDs. This provides a basis for the identification of potential parent material within their wild populations for drought tolerance breeding.

**Highlight:** Wild banana species respond significantly different to water deficit caused by VPD increases and differ in the rate of stomatal reduction, revealing opportunities for drought tolerance breeding.

## 2 Introduction

Climate change projections predict that global temperatures will continue to increase this century (IPCC, 2021). This temperature rise is not only affecting plant functioning directly, but is also increasing air vapour pressure deficit (VPD) (Hatfield and Prueger, 2015; Ficklin and Novick, 2017; Grossiord *et al*., 2020). VPD represents the atmospheric water vapour demand and is defined as the difference between the saturation and actual vapour pressure in the atmosphere (Monteith and Unsworth, 2013). The saturation vapour pressure, the water vapour that air can hold, increases exponentially with temperature and has been increasing as global temperatures rise (Lawrence, 2005). The actual vapour pressure (i.e. absolute humidity in the air) on the other hand has not been rising at the same rate as the saturation vapour pressure, therefore increasing the worldwide VPD (Ficklin and Novick, 2017; Grossiord *et al*., 2020). The impact of this rising VPD is often underestimated compared to other climate change consequences, but periods of high VPD have recently been linked with large-scale tree mortality (Breshears *et al*., 2013; Williams *et al*., 2013) and strong yield reductions (Challinor and Wheeler, 2008; Lobell *et al*., 2013).

Plants respond to the vapour pressure deficit encountered at the leaf level, the leaf-to-air vapour pressure deficit (VPD_leaf_). The leaf temperature can after all deviate from that of the ambient air by transpirational cooling or heating through radiant energy. For a given stomatal opening, transpiration would increase linearly with VPD_leaf_, without any gain in carbon uptake. Stomatal conductance (*g*_*s*_) however decreases with increasing VPD_leaf_, avoiding excessive water loss, but restricting carbon uptake (Dai, Edwards and Ku, 1992; Monteith, 1995; Oren *et al*., 1999). In angiosperms the reduction of *g*_*s*_ in response to an increase in VPD_leaf_ is believed to be abscisic acid (ABA) mediated (Xie *et al*., 2006; Bauer *et al*., 2013; McAdam and Brodribb, 2015). Upon an increase in VPD_leaf_, g_*s*_ is reduced by a rapid ABA biosynthesis (i.e. within 20 min) presumably located in the leaf phloem parenchyma cells and stomatal guard cells (Kuromori, Sugimoto and Shinozaki, 2014; McAdam, Sussmilch and Brodribb, 2016). The trigger for ABA interference under high VPD_leaf_ is believed to be a drop in water status (McAdam and Brodribb, 2016; Sack, John and Buckley, 2018), which has been linked to a limited maximal hydraulic conductance at the leaf, stem and/or root level in comparison to the transpiration (Brodribb and Jordan, 2008; Zhang *et al*., 2013; Choudhary *et al*., 2014; Ocheltree, Nippert and Prasad, 2014; Schoppach *et al*., 2016). Essential gatekeepers for this hydraulic conductance are aquaporins. They are present all along the water transport pathway from root to stomata. Aquaporins were less abundant in soybean and pearl millet genotypes that showed a reduced transpiration rate at high VPD_leaf_ (Sadok and Sinclair, 2010; Devi, Sinclair and Taliercio, 2015; Reddy *et al*., 2017).

Despite the reductions in *g*_*s*_, the transpiration rate usually increases with increasing VPD_leaf_. Only at high VPD_leaf_ significant decreases in transpiration rates have been observed (Franks, Cowan and Farquhar, 1997; Fletcher, Sinclair and Allen, 2007; Gholipoor *et al*., 2010; Ryan *et al*., 2016). These transpiration responses are commonly described by a segmented pattern where the slope of transpiration rate versus VPD_leaf_ is significantly reduced after a specified breakpoint. Significant differences in segmented transpiration responses to VPD_leaf_ have been observed across- and within-species (Fletcher, Sinclair and Allen, 2007; Gholipoor *et al*., 2010; Ryan *et al*., 2016). While some species or genotypes already reduce transpiration rate significantly at low VPD_leaf_, others show only a reduction at higher VPD_leaf_ or even maintain the increasing transpiration rate. Restricting transpiration rate at high VPD has been suggested as a key drought tolerance breeding trait as excessive water loss is avoided and might be saved for later in the growing season (Vadez, 2014; Sinclair *et al*., 2017). Limiting transpiration above a VPD threshold can increase the daily transpiration efficiency but the reduced water use may compromise the yield potential. Reduced transpiration limits carbon uptake, thereby hampering photosynthesis and yield (Richards, 2000; Lee *et al*., 2020; Eyland *et al*., 2021). Moreover, care must be taken that the so-called saved water is not merely lost by evaporation or transpiration by neighbouring plants.

The transpiration rate response to VPD was shown to be highly heritable in wheat (Schoppach *et al*., 2016). Models predict that in drought-prone environments limiting transpiration at high VPD would improve maize and soybean yields by maintaining more soil water available later in the season during flowering or grain filling (Sinclair *et al*., 2010; Messina *et al*., 2015). In these drought-prone regions, the negative effect of *g*_*s*_ reduction on *A* during vegetative growth could be compensated later in the growing season (Sinclair *et al*., 2010; Messina *et al*., 2015). Improved maize hybrids which, amongst other traits, showed reduced transpiration at high VPD_leaf_ indeed increased yields under water-limited conditions (Gaffney *et al*., 2015), while for durum wheat cultivars this was only the case under severe drought conditions (Medina *et al*., 2019).

The current set of edible bananas is complex and has resulted from different parental routes and several back crosses (De Langhe *et al*., 2010; Perrier *et al*., 2011; Martin, Baurens, *et al*., 2020; Cenci *et al*., 2021). The hybrid banana genomes are unbalanced with respect to the parental ones, and inter- and intra-genome translocation chromosomes are relatively common (Christelová *et al*., 2017; Němečková *et al*., 2018). Most, if not all, cultivars have genomes consisting of different proportions of A- and B-genome chromosomes and/or recombinant chromosomes originating from different parents. Similar to other tropical species, bananas are very sensitive to VPD, with reductions in transpiration when VPD exceeds 2 – 2.3 kPa (Aubert and Catsky, 1970; Carr, 2009; Eyland *et al*., 2022). Thomas et al. (1998) observed a diverse response in three banana cultivars with different genomic constitutions. Despite these efforts, the transpiration responses to VPD remain largely uncharacterized across diverse banana genotypes.

The main objective of this work was to evaluate 8 wild banana (sub)species for their stomatal and transpiration responses under increasing VPD at the leaf and whole-plant level. Transpiration rate limitations at high VPD have been indicated as a key breeding trait for high water use efficiency. This work could therefore provide the basis for systematically screening crop wild relatives of banana for their transpiration at high VPD, with the aim to identify potential parent material for drought tolerance breeding.

## 3 Materials & methods

### 3.1 Plant material & growing conditions

A diversity panel of 9 wild banana genotypes belonging to 8 (sub)species (Table 1) were phenotyped for their transpiration response to VPD. Plants were grown in 2.5 L pots filled with peat-based compost and maintained under well-watered conditions. Plants were grown in the greenhouse for 6 - 8 weeks before moving to the growth chamber (Bronson PGC-1400, the Netherlands). The growth chamber contained an air mixing fan and LED panels providing a light intensity of 250 µmol m^-2^ s^-1^ for a 12 h photoperiod and a light spectrum with blue:red:far-red ratio of 1 : 1.5 : 0.15. Plants were acclimated to the growth chamber for one day under a day/night temperature and relative humidity of 27/24.5 °C and 78 %, respectively. The next day the VPD step-changes were initiated by altering relative humidity, while temperature was maintained at 36 °C during this day. VPD was increased by decreasing relative humidity as temperature fluctuations would not only affect VPD but also aquaporin conductance and water viscosity in xylem and mesophyll cells (Matzner and Comstock, 2001; Yang *et al*., 2012). At light onset relative humidity was maintained for 90 min at 87 %, after which it was subsequently decreased to 78, 68, 62 and 56 %, each for 60 min. Average VPDs at each step were 0.77, 1.36, 1.93, 2.34 and 2.64 kPa. Plants were maintained under well-watered conditions by daily watering before light onset. Measurements were taken before 14:00 to avoid afternoon stomatal closure (van Wesemael *et al*., 2019; Eyland *et al*., 2021).

**Table 1:** Wild banana genotypes screened for their transpiration response to increasing vapour pressure deficit (VPD) at both leaf and whole-plant level.

| Name | Subspecies | ITC or collection code<sup>1</sup> | Origin <sup>2</sup> | Collection site<sup>3</sup> | Collection coordinates<sup>3</sup> |
| --- | --- | --- | --- | --- | --- |
| Balbisiana | *Musa balbisiana* | / | Southeast China, northern Indo-Burma, Southwest India, Sri Lanka, Philippines, New Guinea | Japan (Amami) | / |
| Banksii_11 | *Musa acuminata* ssp. *banksii* | SJP416 | New Guinea | Papua New Guinea (Madang) | 5° 37' 8" S<br>145° 28' 7" E |
| Banksii_17 | *Musa acuminata* ssp. *banksii* | SJP814 | New Guinea | Papua New Guinea (Morobe) | 6° 44' 42" S<br>146° 43' 51" E |
| Burmannica | *Musa acuminata* ssp. *burmannica* | ITC0283 | southern Indo-Burma | / | / |
| Burmannicoides | *Musa acuminata* ssp. *burmannicoides* | ITC0249 | southern Indo-Burma | / | / |
| Errans | *Musa acuminata* ssp. *errans* | ITC1028 | Philippines | / | / |
| Malaccensis_33 | *Musa acuminata* ssp. *malaccensis* | 928533 | Sumatra and Malayan Peninsula | Malaysia (Pahang) | 3°53'51" N<br>102°12'23"E |
| Microcarpa | *Musa acuminata* ssp. *microcarpa* | ITC0253 | Borneo | / | / |
| Zebrina | *Musa acuminata* ssp. *zebrina* | ITC1177 | Sumatra and Malayan Peninsula | / | / |

<sup>1</sup>Genotypes without ITC code were collected germplasm and not yet available at the International Transit Centre (ITC) collection. The collection code represents the given code to the mother plant during collection. <sup>2</sup>Genotype origin as described by Janssens et al. (2016). <sup>3</sup>Only locations of collected samples are shown.

### 3.2 Leaf gas exchange measurements

Gas exchange responses to step increases in VPD_leaf_ were measured every 60 s on the middle of the second youngest fully developed leaf using a LI-6800 infrared gas analyser (LI-COR, USA). Light intensity and CO_2_ concentration were maintained at 250 µmol m^-2^ s^-1^ and 400 µmol mol^-1^, respectively. Leaf temperature was maintained at 36 °C. Relative humidity went from 85 to 75, 65, 55, 45 and 35 %, reaching VPD_leaf_ of 0.91, 1.50, 2.09, 2.69, 3.28 and 3.87 kPa. Note that measurements were stopped early if the drying capacity of the infra-red gas exchange system was saturated and unable to maintain reduced relative humidity. The intrinsic water use efficiency (_i_WUE) was calculated as _i_WUE = *A*/*g*_*s*_ with *A* being the photosynthetic rate. At every VPD_leaf_ level the steady-state *g*_*s*_, *A*, E_rate_ (transpiration rate) and _i_WUE after 60 min was calculated. The maximum *g*_*s*_ was calculated as the highest *g*_*s*_ observed across all VPD_leaf_ levels. Segmented regression was performed on the transpiration rate response to increasing VPD_leaf_ for each genotype by using a nonlinear mixed effect model in which the intercept was assumed to vary at individual plant level (segmented R package, Muggeo, 2008). This analysis calculates the optimal breakpoint in the transpiration response with a different linear response before and after the breakpoint. To determine the effect of the reduction in stomatal opening on the transpiration, the transpiration reduction (ϕ_E_) was determined according to Franks *et al*. (1999) and Ryan *et al*. (2016) (Fig. 1). For each individual, a linear regression was fitted through the transpiration rate at the first two VPD_leaf_ levels (0.90 and 1.50 kPa). This linear regression was then extrapolated to predict the transpiration rate (E_pred_) at higher VPD_leaf_ levels (2.69, 3.28 and 3.87 kPa) (Fig. 1). The percentage decrease of the actual measured transpiration rate (E_meas_) compared to E_pred_ (Fig. 1) was then quantified at each VPD_leaf_ level:

**Fig. 1:**
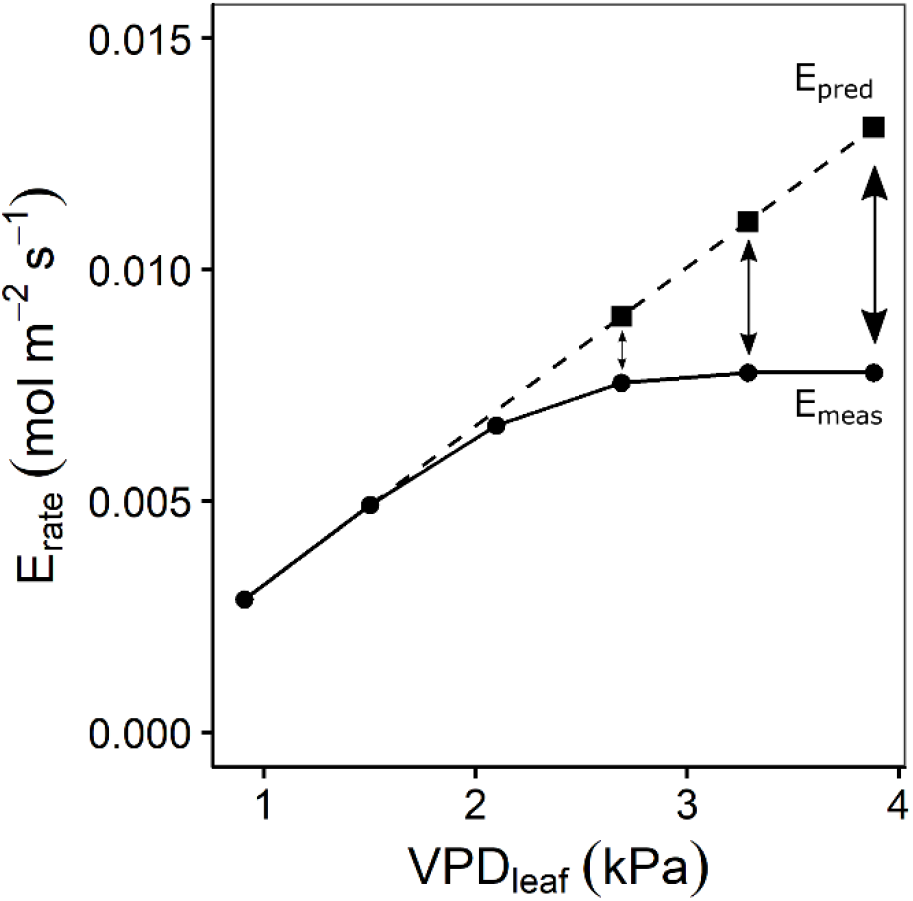
Quantification of the transpiration reduction (ϕ_E_) according to Franks et al. (1999) and Ryan et al. (2016). A linear regression was fitted through the transpiration rate at the first two air-to-leaf vapour pressure deficit (VPD_leaf_) levels. This linear regression was extrapolated (dashed line) to estimate the transpiration rate (E_pred_) at VPD_leaf_ of 2.69, 3.28 and 3.87 kPa. E_pred_ was then compared to the measured transpiration rate (E_meas_) to calculate ϕ_E_ (Eq. 1).

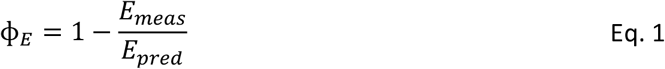

The percentage of limitation of the photosynthetic rate (*A*) by *g*_*s*_ reduction was calculated at every VPD_leaf_ level by comparing the measured *A* (*A*_meas_) with the overall maximally measured *A* (*A*_max_):

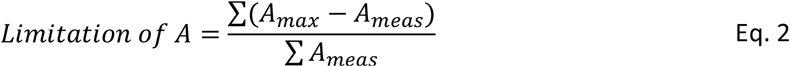

Stomatal reduction (ϕ_stom_) with increasing VPD was defined as the absolute slope between stomatal conductance (*g*_*s*_) and log_e_(VPD_leaf_) as described by Oren *et al*. (1999):

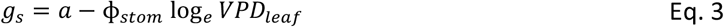

where *a* is the estimated *g*_*s*_ at VPD_leaf_ 1 kPa.

### 3.3 Whole-plant transpiration rate

Plants were placed on balances (0.01 g accuracy, Kern, Germany) to register their weight every 10 s. The soil was covered by plastic to avoid evaporation and ensure only water loss through transpiration. Transpiration during each VPD step was calculated by differentiating 5 min average total weight (m_*tot*_) at the start of the VPD level with the 5 min average total weight at the end of the VPD level:

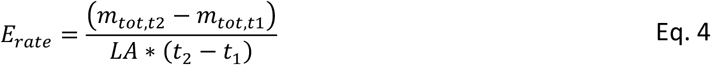

Transpiration was normalized by leaf area (LA) and the time (t) passed. LA was quantified by destructive leaf area imaging at the end of the experiment.

Segmented regression was performed on the transpiration rate response to increasing VPD for each genotype by using a nonlinear mixed effect model in which the intercept was assumed to vary at plant level. Transpiration reduction (ϕ_E_) was determined according to Eq. 1 with linear regression between the two first VPD levels (0.77 and 1.36 kPa) and comparison between E_pred_ and E_meas_ at the highest level (2.64 kPa).

### 3.4 Statistics

All data processing and statistical analysis were carried out in R (V3.6.2). Genotypic differences were tested by applying analysis of variance (ANOVA) with a post hoc Benjamini & Hochberg correction. Significance of the segmented response of transpiration rate to VPD compared to a linear response was determined by the Davies Test (segmented R package, Muggeo, 2008). K-means clustering of genotypes was performed on the average scaled output of the segmented regression, the transpiration reduction, the stomatal reduction and photosynthesis limitation, including measurements by leaf gas exchange and by whole-plant transpiration were included (Hartigan and Wong, 1979). Clusters were optimized across 10,000 random sets of cluster centres and plotted on the first two principal components.

## 4 Results

### 4.1 Diverse response to VPD: three phenotypic clusters

The transpiration response was measured at leaf and whole plant level while relative humidity was stepwise decreased and VPDs consequently increased. The response to increasing VPD at leaf and whole-plant level was described by the segmented regression of transpiration rate versus VPD, the transpiration reduction (Eq. 1), the photosynthetic limitation under increasing VPD (Eq. 2) and the stomatal reduction (Eq. 3). K-means clustering was performed on the output variables measured by both leaf gas exchange and whole-plant transpiration (Table 2). Three clusters were identified and plotted along the first two principal components (Fig. 2). The first principal component was mainly determined by the limitation of photosynthetic rate (*A*) at high VPDs and the transpiration reduction at leaf and whole-plant level (Table 2). Important variables in the second principal component were the slope before the breakpoint in transpiration rate with increasing VPD and the stomatal reduction (Table 2). Cluster I consisted of only one genotype: *M. acuminata* ssp. *errans* (Fig. 2). In group II *M. acuminata* ssp. *zebrina* and *M. balbisiana* clustered together (Fig. 2). Group III contained 6 genotypes: *M. acuminata* ssp. *banksii*, ssp. *burmannica*, ssp. *burmannicoides*, ssp. *malaccensis* and ssp. *microcrocarpa* (Fig. 2).

**Table 2:** Variables included in the k-means clustering and their principal component (PC) loadings.

| Variable<sup>1</sup> | Measurement level | PC1 loading<sup>2</sup> | PC2 loading |
| --- | --- | --- | --- |
| Limitation of A at 3.87 kPa | Leaf gas exchange | -0.35 | 0.08 |
| Limitation of A at 3.28 kPa | Leaf gas exchange | -0.34 | 0.13 |
| Transpiration reduction at 2.69 kPa | Leaf gas exchange | -0.34 | -0.14 |
| Limitation of A at 2.69 kPa | Leaf gas exchange | -0.32 | 0.23 |
| Transpiration reduction at 3.28 kPa | Leaf gas exchange | -0.32 | -0.24 |
| Transpiration reduction at 3.87 kPa | Leaf gas exchange | -0.29 | -0.34 |
| Transpiration reduction at 2.64 kPa | Whole plant transpiration | -0.29 | 0.00 |
| Breakpoint in transpiration rate | Leaf gas exchange | 0.25 | -0.04 |
| Slope after breakpoint in transpiration rate | Whole plant transpiration | 0.24 | 0.19 |
| Limitation of A at 2.09 kPa | Leaf gas exchange | -0.22 | 0.22 |
| Slope after breakpoint in transpiration rate | Leaf gas exchange | 0.20 | 0.28 |
| Slope before breakpoint in transpiration rate | Leaf gas exchange | 0.16 | -0.39 |
| Breakpoint in transpiration rate | Whole plant transpiration | 0.14 | 0.05 |
| Limitation of A at 1.50 kPa | Leaf gas exchange | 0.13 | -0.03 |
| Slope before breakpoint in transpiration rate | Whole plant transpiration | 0.09 | -0.49 |
| Stomatal reduction | Leaf gas exchange | 0.04 | -0.41 |

<sup>1</sup>Variables measured by leaf gas exchange and at whole-plant transpiration were included. <sup>2</sup>Data were ordered following the absolute value of the first principal component loadings.

**Fig. 2:**
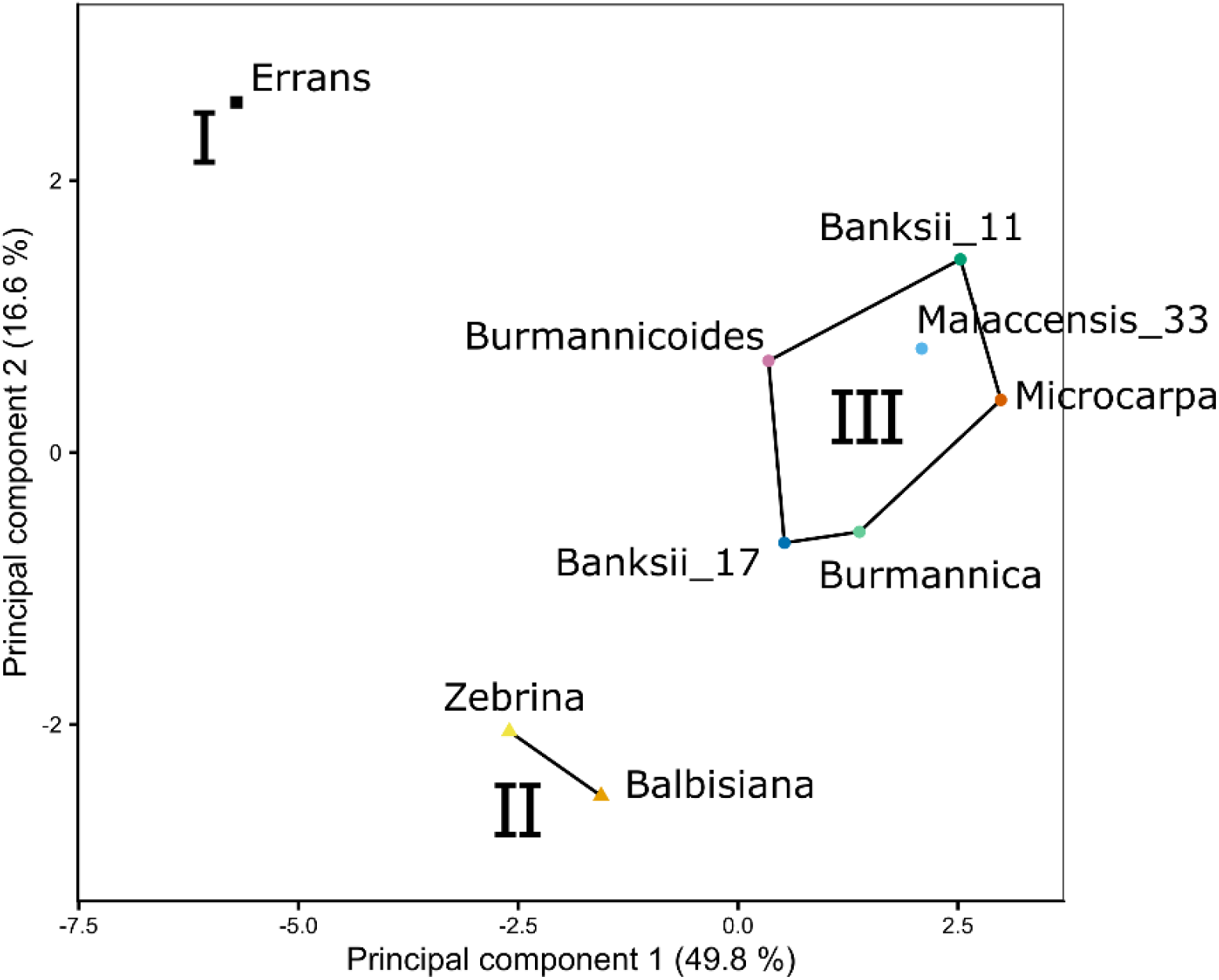
Three genotype groups (I, II, III) were defined by k-means clustering based on the stomatal reduction, transpiration reduction and photosynthetic limitation under increasing VPD (see variables in Table 2). Both variables measured by leaf gas exchange and at whole-plant transpiration were included. Lines and regions represent the three genotype groups from k-means clustering plotted along the first two principal components (Table 2). The first principal component was mainly determined by the limitation of photosynthetic rate at high VPDs and the transpiration reduction at leaf and whole-plant level. Important variables in the second principal component were the slope before the breakpoint in transpiration rate with increasing VPD and the stomatal reduction.

### 4.2 Leaf level responses of *g*_*s*_, transpiration rate and *A* to increasing VPD_leaf_

With increasing VPD_leaf_, *g*_*s*_ decreased in all genotypes (Fig. 3A, Supplemental Table S1). The transpiration rate initially increased, but eventually reached steady-state or even declined (Fig. 3B). The transpiration rate and *g*_*s*_ of *M. acuminata* ssp. *errans* were lowest and differed significantly from all other genotypes at VPD_leaf_ exceeding 1.50 and 2.09 kPa, respectively (Fig. 3A-B, Supplemental Table S1). Under a VPD_leaf_ ≤ 2.9 kPa, the highest transpiration rates and *g*_*s*_ were observed for *M. balbisiana* and *M. acuminata* ssp. *burmannica*. However, when VPD_leaf_ increased further, the *g*_*s*_ of *M. balbisiana* decreased stronger than *M. acuminata* ssp. *burmannica*, translating only in *M. balbisiana* in a lower transpiration rate (Fig. 3A-B, Supplemental Table S1). As *g*_*s*_ decreased with increasing VPD_leaf_, the CO_2_ uptake was limited and *A* decreased (Fig. 3C). The lowest *A* was observed for *M. acuminata* ssp. *errans* and ssp. *burmannicoides*, with significantly lower *A* compared to all other genotypes except *M. acuminata* ssp. *zebrina* (Fig. 3C, Supplemental Table S1). The intrinsic water use efficiency (_i_WUE) increased with increasing VPD_leaf_ (Fig. 3D). _i_WUE was highest in *M. acuminata* ssp. *errans* and differed significantly from all other genotypes as VPD_leaf_ exceeded 1.5 kPa (Fig. 3D, Supplemental Table S4.1). The lowest _i_WUE were observed for *M. acuminata* ssp. *burmannica* and ssp. *burmannicoides* (Fig. 3D).

**Fig. 3:**
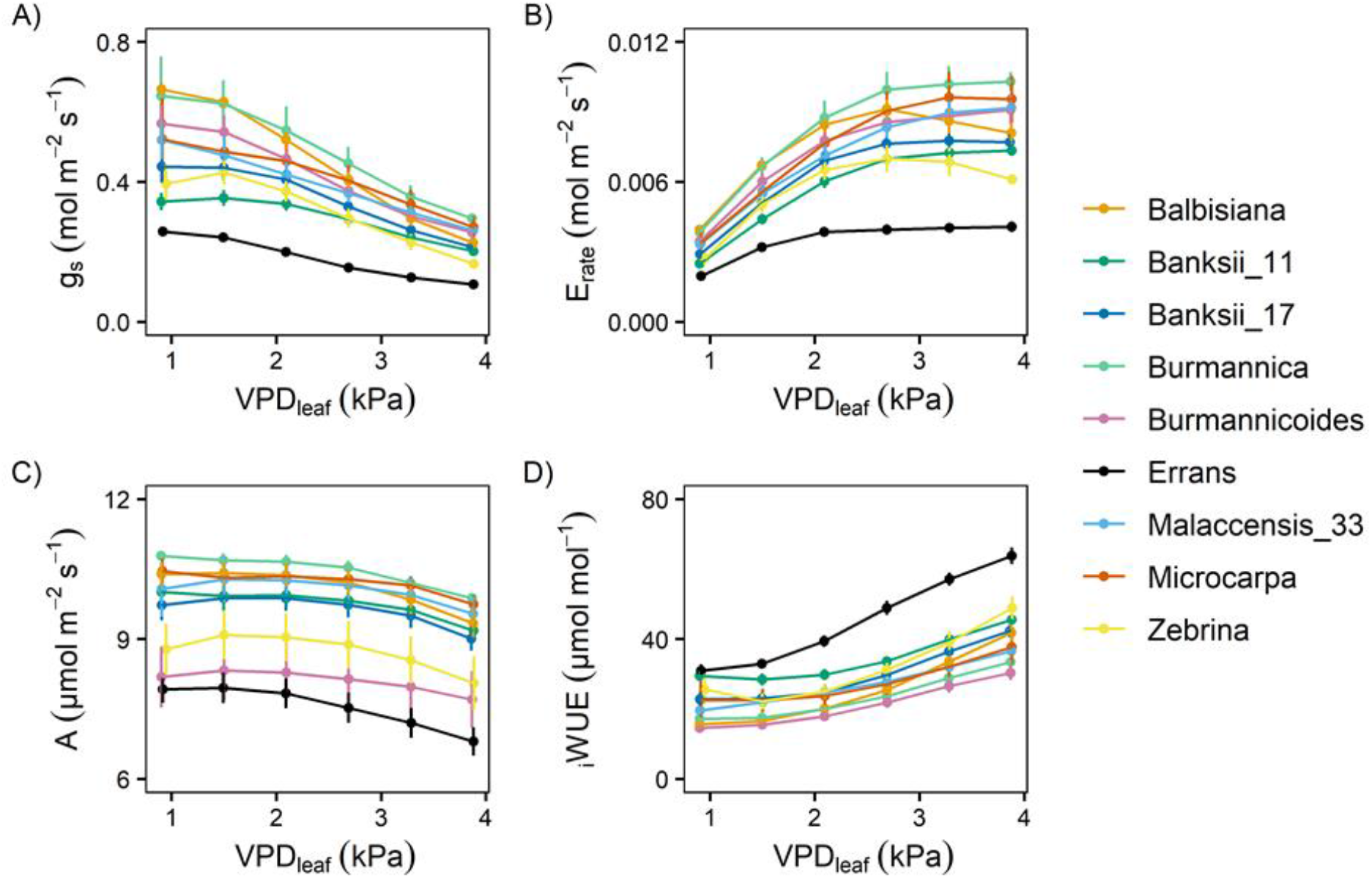
Gas exchange response to step-increases in leaf-to-air vapour pressure deficit (VPD_leaf_) for 9 wild banana genotypes. Steady-state response of (A) stomatal conductance (*g*_*s*_), (B) transpiration rate (E_rate_), (C) photosynthetic rate (*A*) and (D) intrinsic water use efficiency (_i_WUE) to increasing VPD_leaf_. Data represent mean±se values after 60 min at a specific VPD_leaf_ level (n=3-7). Significance is shown in Supplemental Table S1.

In all genotypes there was a decrease in the slope of transpiration rate versus VPD_leaf_ (Fig. 3B). This response was described by a segmented regression with a specified breakpoint after which the slope of the transpiration rate decreases. A significant breakpoint in transpiration rate in response to VPD_leaf_ was identified in all genotypes (Fig. 4). Across genotypes the breakpoints ranged between 1.75 and 2.5 kPa with *M. acuminata* ssp. *errans* having a significant breakpoint at the lowest VPD_leaf_ (Fig. 4, Fig. 5). Two *M. acuminata* ssp. *banksii* genotypes and ssp. *microcarpa* showed the highest breakpoint in transpiration rate (Fig. 4, Fig. 5). The groups defined by k-means clustering differed in their segmented transpiration response (Fig. 5). Group I consisted only of *M. acuminata* ssp. *errans*, the genotype with a breakpoint (a reduction in transpiration rate) at the lowest VPD_leaf_, as well as the lowest slope (the lowest E_rate_) before the breakpoint (Fig. 5). Group II, consisting of *M. acuminata* ssp. *zebrina* and *M. balbisiana*, had a breakpoint at a relatively low VPD_leaf_ around 2 kPa and a negative slope after the breakpoint (Fig. 5). This negative slope indicates a net decrease in transpiration rate, which was not observed in the other genotypes. In group III all genotypes kept relatively high transpiration rates at relatively high VPD_leaf_. *Musa acuminata* ssp. *burmannica*, ssp. *burmannicoides* and ssp. *malaccensis* had a breakpoint at relatively low VPD_leaf_, but maintained a high slope of transpiration rate afterwards while the *M. acuminata* ssp. *banksii* genotypes and ssp. *microcarpa* showed only a significant breakpoint in transpiration rate at higher VPD_leaf_, (Fig. 5).

**Fig. 4:**
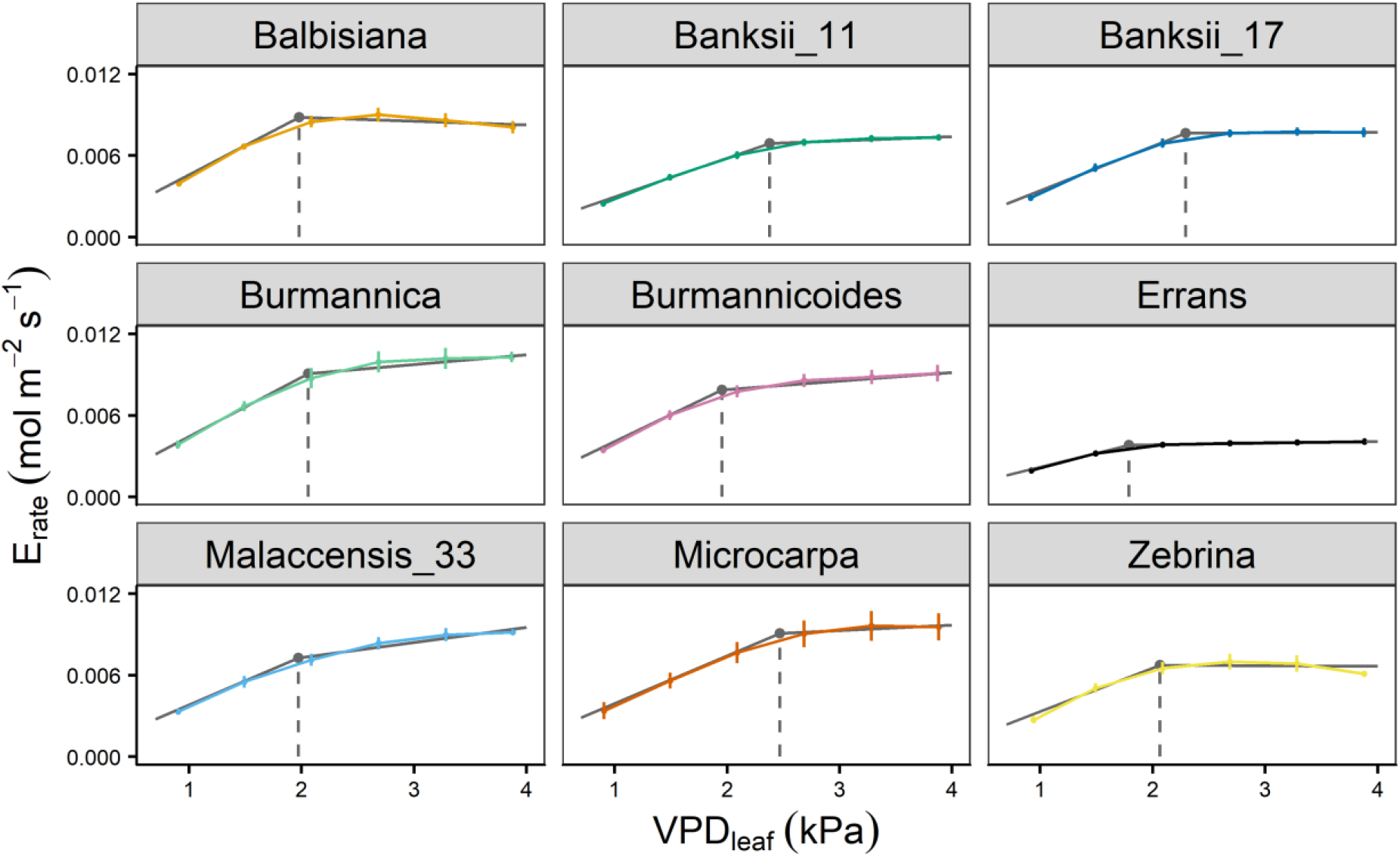
Transpiration rate response of 9 wild banana genotypes to step-increases in leaf-to-air vapour pressure deficit (VPD_leaf_). A significant breakpoint in transpiration rate was identified for all genotypes (P-value Davies Test < 0.05). Solid grey lines represent slopes of the modelled segmented response. Grey point and dashed grey line represent the breakpoint in transpiration rate and the VPD_leaf_ of the breakpoint. Data represent mean±se values after 60 min at a specific VPD_leaf_ level (n=3-7).

**Fig. 5:**
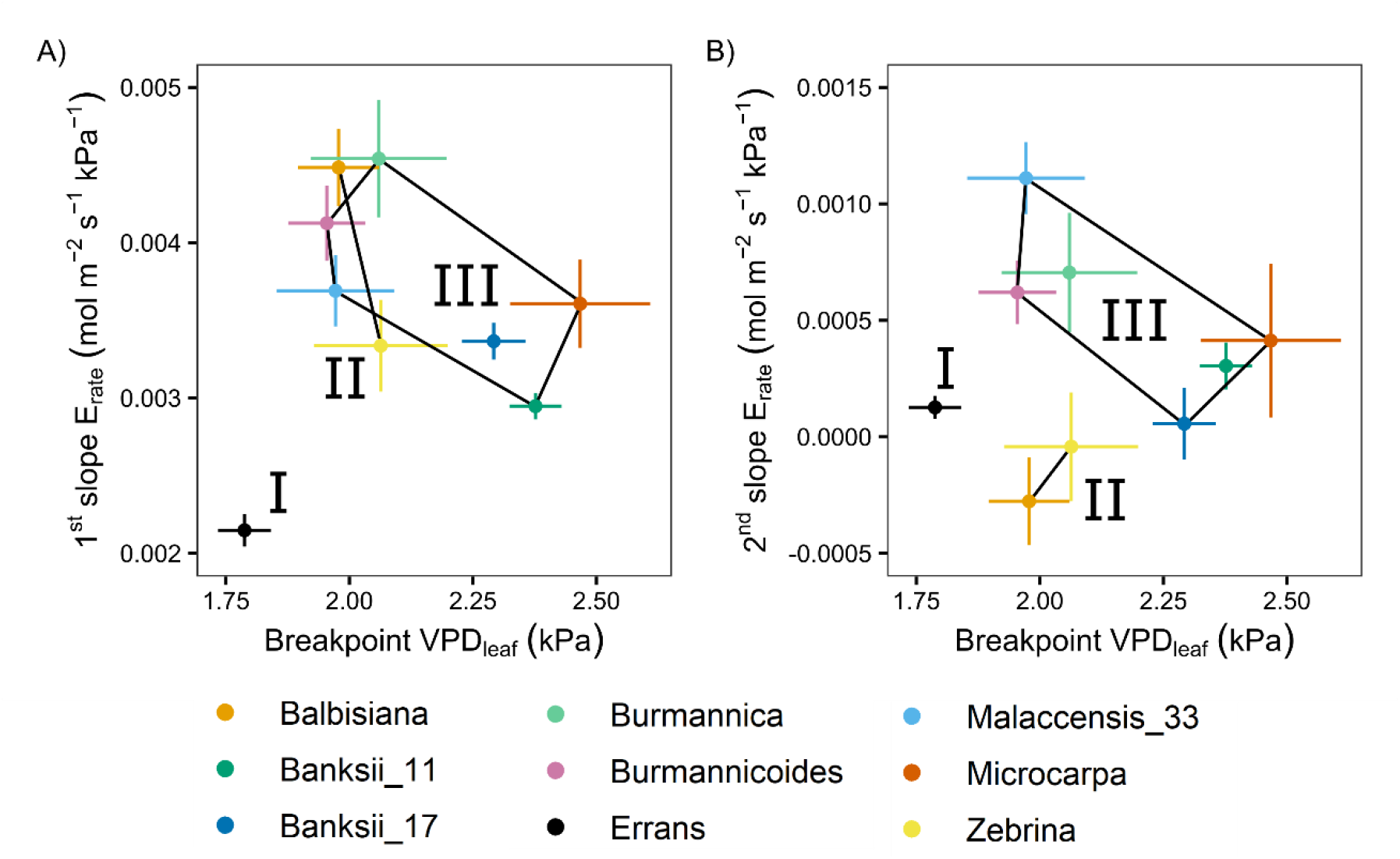
Slopes and breakpoints of the segmented transpiration rate response to step-increases in leaf-to-air vapour pressure deficit (VPD_leaf_). (A) Relation between the breakpoint in transpiration rate and the slope before the breakpoint. (B) Relation between the breakpoint in transpiration rate and the slope after the breakpoint. Three groups (I, II, III) were defined by k-means clustering and are represented by black lines connecting the included genotypes. All segmented responses were significant (P < 0.05). Data represent the optimal estimated value±se. (n=3-7).

The transpiration reduction (ϕ_E_) (Eq. 1, Fig. 1) representing the increase in stomatal resistance with increasing VPD_leaf_ also differed significantly across genotypes (Fig. 6A, Supplemental Table S2). Reductions in transpiration ranged between 37 and 59 % at the highest VPD_leaf_ of 3.87 kPa (Fig. 6A, Supplemental Table S2). The highest reductions in transpiration were observed for *M. acuminata* ssp. *errans*, ssp. *zebrina* and *M. balbisiana* (Fig. 6A). The transpiration reduction of group I and II was significantly higher compared to group III at all VPD_leaf_ levels (Fig. 6A, Supplemental Table S2).

**Fig. 6:**
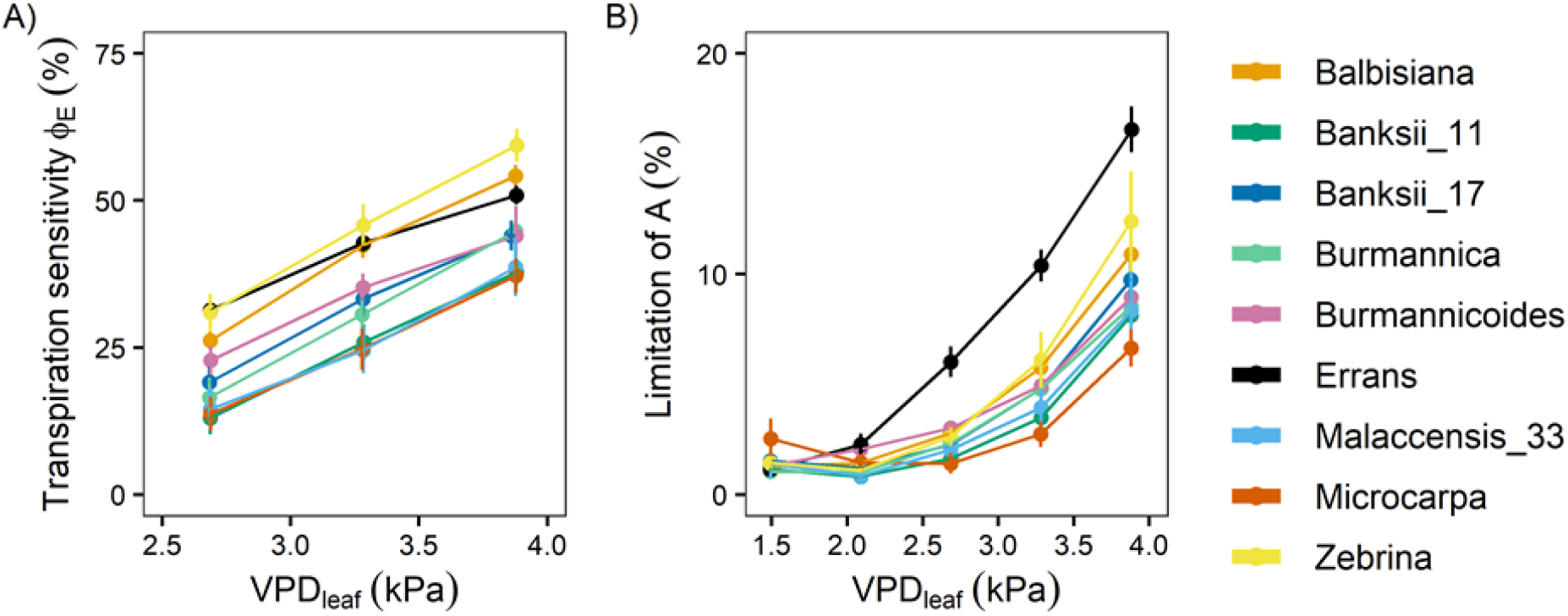
Transpiration reduction (ϕ_E_) and limitation of photosynthetic rate (*A*) with increasing leaf-to-air vapour pressure deficit (VPD_leaf_). (A) ϕ_E_ in response to increasing VPD_leaf_. ϕ_E_ was determined as shown in Eq. 1. (B) Limitation of *A* in response to increasing VPD_leaf_. The limitation of *A* was determined as shown in Eq. 2. Data represent mean±se. (n=3-7). Significance in shown in Supplemental Tables S2 and S3.

The decrease in stomatal opening with increasing VPD_leaf_ limited the photosynthetic rate (*A*). In all genotypes there was a significant increase in the limitation of *A* with increasing VPD_leaf_ (P < 0.01) and the limitation ranged from 7 to 17 % at the highest VPD_leaf_ level (Fig. 6B, Supplemental Table S3). The limitation of *A* was highest in *M. acuminata* ssp. *errans* from VPD_leaf_ 2.69 kPa onwards, followed by *M. acuminata* ssp. *zebrina* and *M. balbisiana* (Fig. 6B, Supplemental Table S3). The limitation of *A* was significantly higher in group I compared to group II and III from VPD_leaf_ 2.69 kPa onwards (Supplemental Table S3). At VPD_leaf_ of 3.28 and 3.87 kPa group II had a significantly higher *A* limitation compared to group III (Supplemental Table S3). Across genotypes the limitation of *A* at higher VPD_leaf_ (≥ 2.69 kPa) was significantly correlated to the breakpoint in transpiration rate (R^2^ = 0.47-0.57; Supplemental Fig. S1). Similarly, the limitation of *A* and the transpiration reduction at higher VPD_leaf_ (≥ 2.69 kPa) were significantly correlated (R^2^ = 0.53-0.58; Supplemental Fig. S1). These correlations indicate that strong reductions in transpiration at high VPD_leaf_ result in higher *A* limitations.

The stomatal reduction (ϕ_stom_), defined as the slope of *g*_*s*_ versus log_e_(VPD_leaf_) (Eq. 3) differed significantly across genotypes (Supplemental Table S4). Highest stomatal reduction was observed in *M. balbisiana*, while *M. acuminata* ssp. *errans* showed lowest reduction (Fig. 7, Supplemental Table S4). The stomatal reduction was strongly correlated to the maximum observed *g*_*s*_ (R^2^ = 0.88, Fig. 7, Supplemental Fig. S1). No significant differences across previously described groups was observed (Supplemental Table S4).

**Fig. 7:**
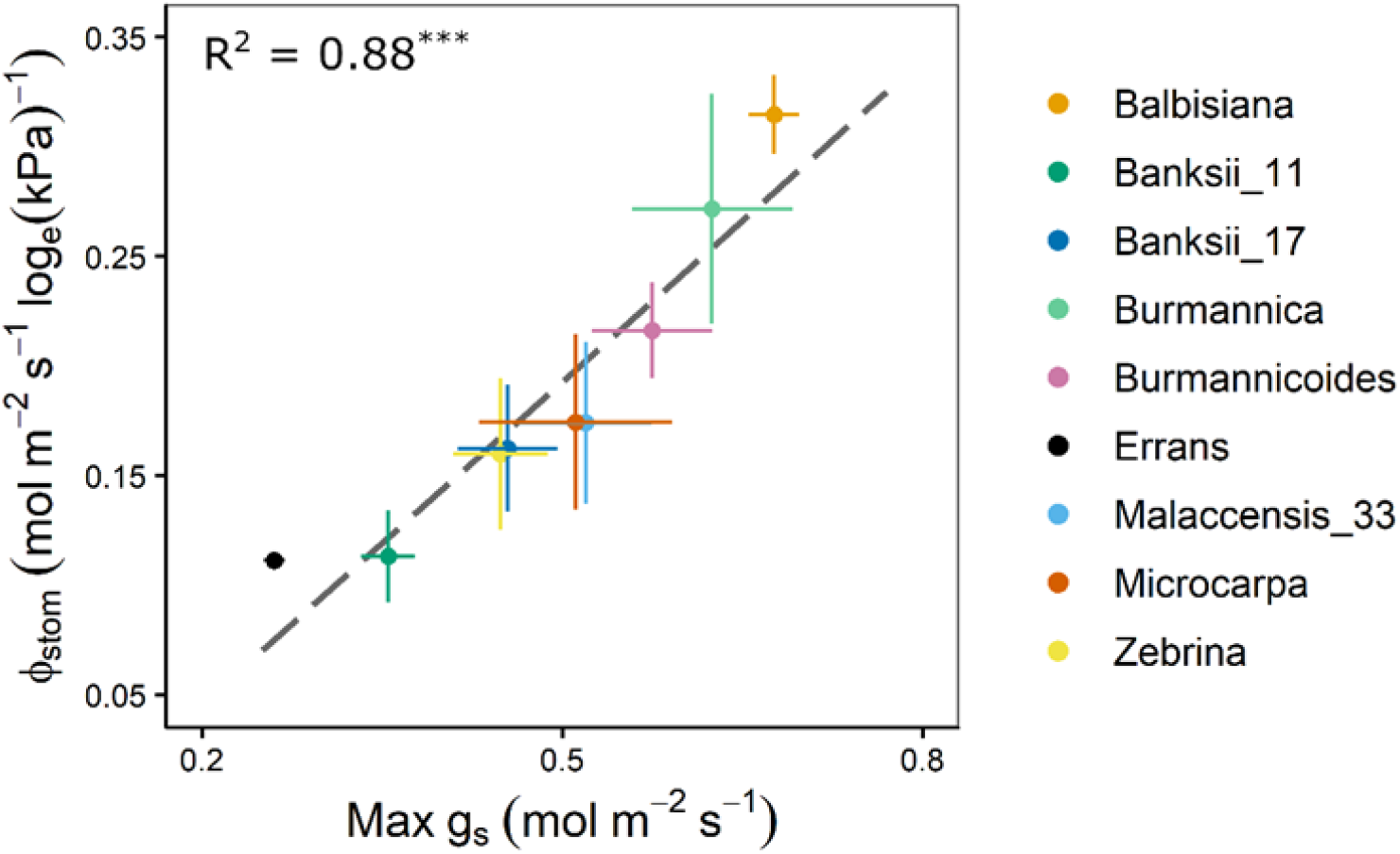
Stomatal reduction (ϕ_stom_) in relation to the maximum observed stomatal conductance (max g_s_). The ϕ_stom_ and max g_s_ were significantly correlated (R^2^ = 0.88, P < 0.001). Data represent mean±se (n=3-7). Significance is shown in Supplemental Table S4.

### 4.3 Whole-plant transpiration rate responses corroborate leaf measurements

The whole-plant transpiration rate increased between 98 and 197 % with increasing VPD (Fig. 8). The lowest transpiration rates were observed for *M. acuminata* ssp. *errans* with significant differences compared to all other genotypes from VPD 1.93 kPa and beyond (Fig. 8, Supplemental Table S5). Transpiration rates of all other genotypes were double compared to *M. acuminata* ssp. *errans* at the highest VPD level (Fig. 8, Supplemental Table S5).

**Fig. 8:**
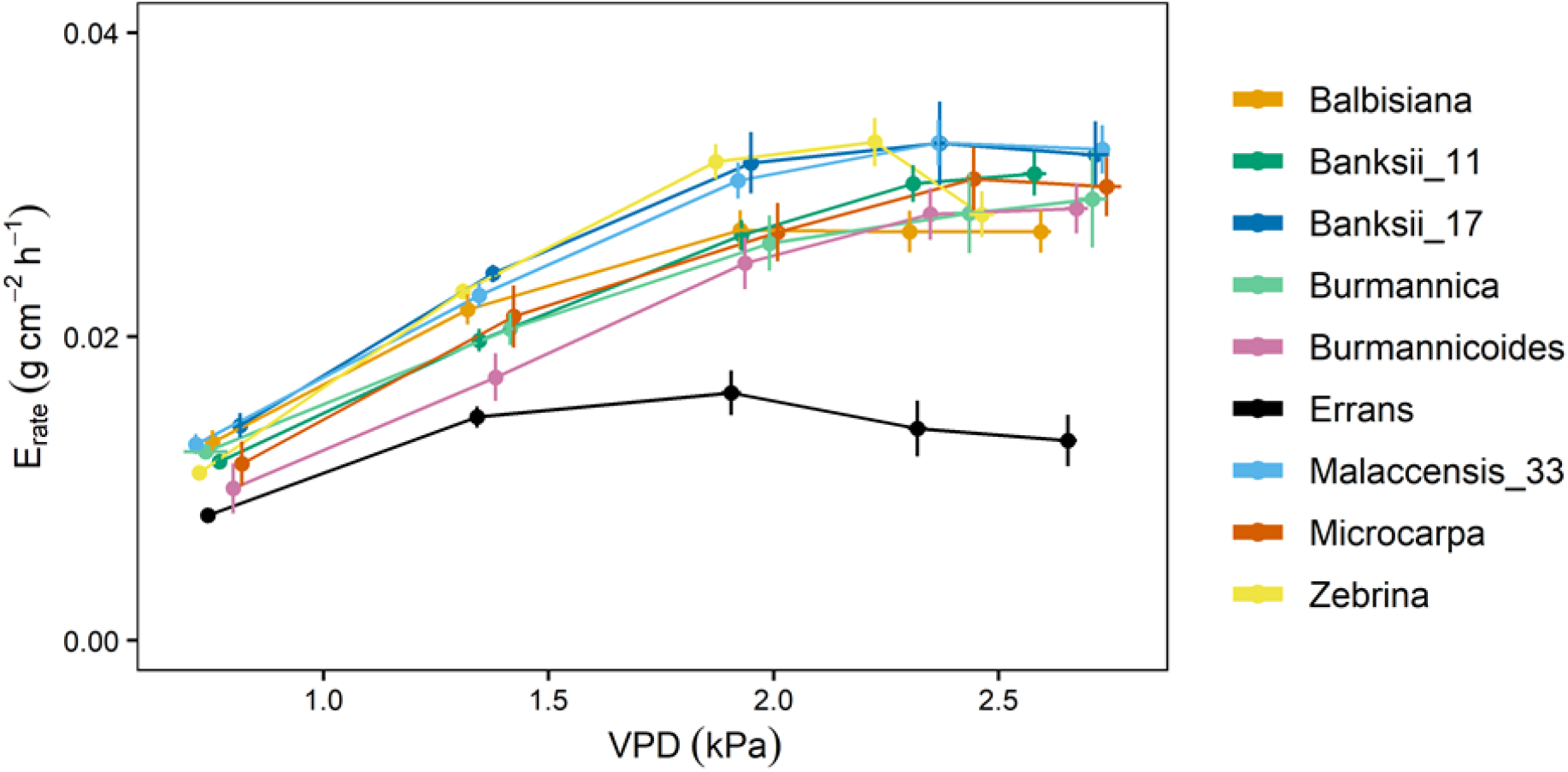
Whole-plant transpiration rate (E_rate_) response to step-increases in air vapour pressure deficit (VPD) for 9 wild banana genotypes. Note that VPD values slightly differed between genotypes depending on the maximal drying capacity of the growth chamber. Data represent mean±se values after 60 min at a specific VPD level (n=4-8). Significance is shown in Supplemental Table S5.

A significant breakpoint in whole-plant transpiration rate response to VPD was identified in all genotypes (Fig. 9). The breakpoints ranged between 1.6 and 2.2 kPa, with *M. acuminata* ssp. *errans* and *M. balbisiana* having the lowest breakpoint (Fig. 9, Fig. 10). The slope after the breakpoint was strongly negative in *M. acuminata* ssp. *errans* and ssp. *zebrina* (Fig. 9, Fig. 10). Genotypes belonging to group I or II thus showed breakpoints in transpiration rate at lower VPD values and/or strongly negative second slopes (Fig. 10).

**Fig. 9:**
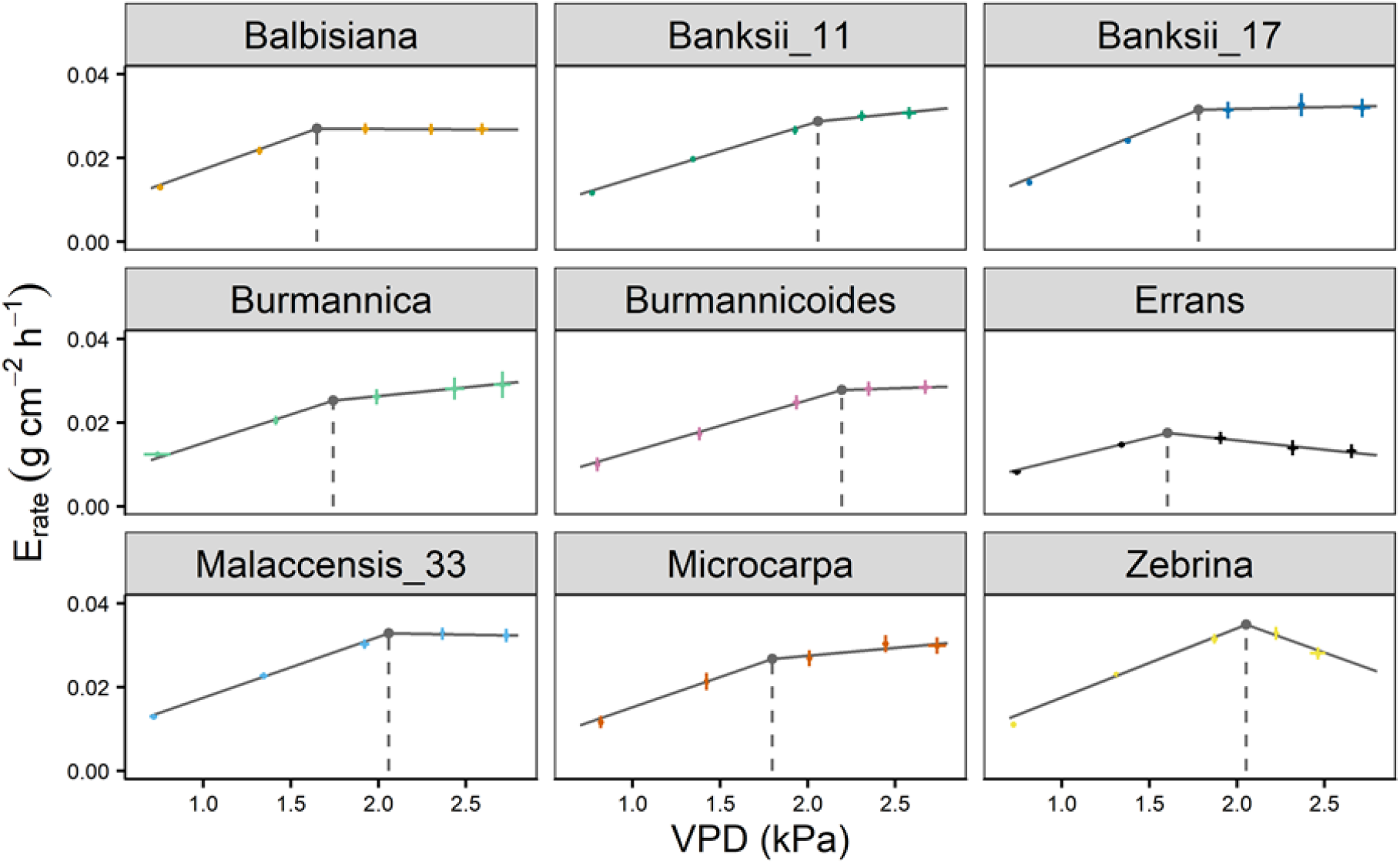
Whole-plant transpiration rate (E_rate_) response of 9 wild banana genotypes to step-increases in air vapour pressure deficit (VPD). A significant breakpoint in transpiration rate was identified for all genotypes (P-value Davies Test < 0.05). Solid grey lines represent slopes of the modelled segmented response. Grey point and dashed grey line represent the breakpoint in transpiration rate and the VPD of the breakpoint. Data represent mean±se (n=4-8).

**Fig. 10:**
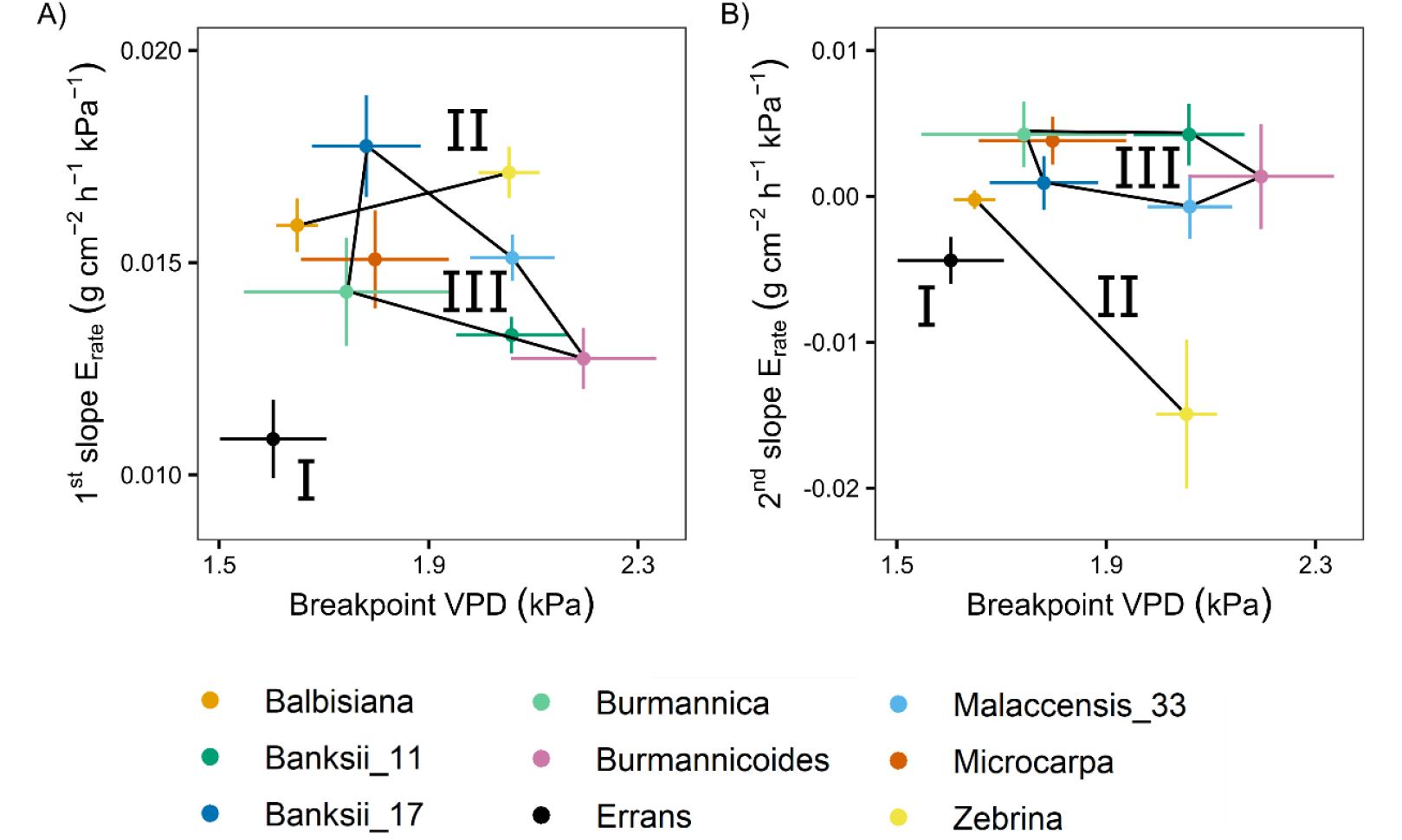
Slopes and breakpoints of the segmented whole-plant transpiration rate (E_rate_) response to step-increases in air vapour pressure deficit (VPD). (A) Relation between the breakpoint in whole-plant transpiration rate and the slope before the breakpoint. (B) Relation between the breakpoint in whole-plant transpiration rate and the slope after the breakpoint. Three groups (I, II, III) were defined by k-means clustering and are represented by black lines connecting the included genotypes. All segmented responses were significant (P < 0.05). Data represent the optimal estimated value± se (n = 4-8).

The whole-plant transpiration reduction (ϕ_E_) (Eq. 1, Fig. 1) of *M. acuminata* ssp. *errans* was significantly higher compared to all other genotypes (Fig. 11, Supplemental Table S6). The second highest transpiration reduction was observed for *M. acuminata* ssp. *zebrina* and *M. balbisiana* (Fig. 11, Supplemental Table S6). Group I (*M. acuminata* ssp. *errans*) showed a significantly higher transpiration reduction compared to group II and III (Supplemental Table S6). Group II (*M. acuminata* ssp. *zebrina* and *M. balbisiana*) showed a significantly higher transpiration reduction compared to group III (*Musa acuminata* ssp. *burmannica*, ssp. *burmannicoides*, ssp. *malaccensis*, ssp. *banksii* and ssp. *microcarpa*) (Supplemental Table S6).

**Fig. 11:**
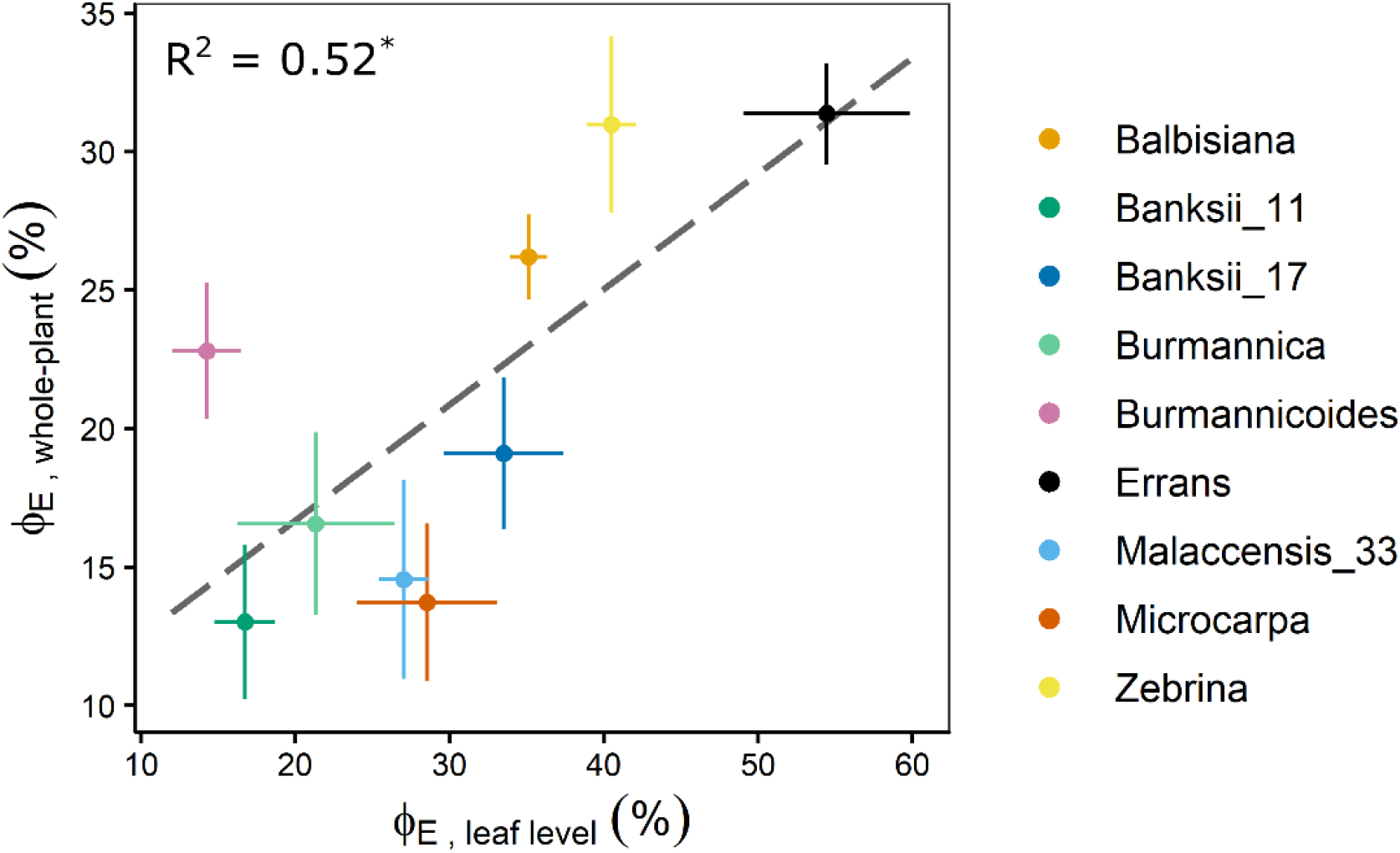
Transpiration reduction measured at whole-plant level (ϕ_E, whole-plant_) at VPD 2.64 kPa in relation to the transpiration reduction measured at leaf level (ϕ_E, leaf level_) at VPD_leaf_ 2.69 kPa. The ϕ_E_ at leaf and whole-plant level were significantly correlated (R^2^ = 0.52, P < 0.05). Data represent mean±se (n=4-8). Significant differences between genotypes or groups are indicated in Supplemental Tables S2 and S6.

The whole-plant transpiration reduction was significantly correlated to the transpiration reduction measured at leaf level at similar VPD (R^2^ = 0.52, Fig. 11, Supplemental Fig. S1). Similarly, the whole-plant transpiration reduction was significantly correlated to the limitation of *A* measured at leaf level for VPD_leaf_ exceeding 2.1 kPa (R^2^ = 0.50 - 0.73, Supplemental Fig. S1).

## 5 Discussion

Diversity in transpiration patterns with increasing VPD has been observed among different genotypes of many crops including chickpea, maize, peanut, pearl millet, sorghum and soybean (Fletcher, Sinclair and Allen, 2007; Gholipoor *et al*., 2010; Jyostna Devi, Sinclair and Vadez, 2010; Kholová *et al*., 2010; Yang *et al*., 2012; Ryan *et al*., 2016; Sivasakthi *et al*., 2017). We observed a significant change in the transpiration rate of 9 wild banana genotypes already at VPD levels between 1.6 and 2.5 kPa (Fig. 4, Fig. 9). These values are in line with the general transpiration rate reduction of banana at VPD 2 to 2.3 kPa reported by Carr (2009) and the modelled VPD responses of (Eyland *et al*., 2022). The breakpoints in transpiration rate were at similar VPDs compared to other crops (Gholipoor *et al*., 2010; Yang *et al*., 2012; Ryan *et al*., 2016). However, in other crops several genotypes were identified without a breakpoint as they maintained a linear increase in transpiration rate with increasing VPD (Fletcher, Sinclair and Allen, 2007; Gholipoor *et al*., 2010; Jyostna Devi, Sinclair and Vadez, 2010; Kholová *et al*., 2010; Yang *et al*., 2012; Ryan *et al*., 2016; Sivasakthi *et al*., 2017). Moreover, temperature and other environmental factors like radiation and soil water potential have been shown to interact with VPD in banana (Eyland *et al*., 2022). These complex interactions explain why a fixed VPD level per genotype, where a reduction in transpiration takes place, cannot be defined without taking the other environmental conditions in account.

The wild banana genotypes clustered in three groups based on their leaf gas exchange and whole-plant transpiration response to VPD (Fig. 2). Genotypes of group I and II, *M. acuminata* ssp. *errans, M. acuminata* ssp. *zebrina* and *M. balbisiana*, showed the highest transpiration rate limitations. This is in line with our previous observations under fluctuating conditions: *M. balbisiana* showed together with *M. acuminata ssp. errans* the most pronounced response by strongly decreasing their transpiration rate (Eyland *et al*., 2022). As reported by Oren *et al*. (1999), the stomatal reduction was significantly correlated to the maximum g_*s*_ (Fig. 7, Supplemental Fig. 1). This indicates that genotypes with higher g_s_ under low VPD_leaf_ show higher stomatal closure at increasing VPD_leaf_. However, *M. acuminata* ssp. *errans* (group I) showed a very strong stomatal response, despite its low g_*s*_. As a consequence of this strong stomatal restriction, the _i_WUE of *M. acuminata* ssp. *errans* was significantly higher compared to all other genotypes (Fig. 3D). In contrast to the very conservative behaviour of *M. acuminata* ssp. *errans*, the genotypes of group II displayed high *g*_*s*_ and *A* when VPD_leaf_ was favourable in addition to early or strong transpiration rate reductions at high VPD_leaf_. This behaviour is assumed to be beneficial in drought-prone areas with periods of high VPD (Sadok and Sinclair, 2010; Vadez, 2014), as water is used efficiently and saved for later in the growing season. Some genotypes of group III also showed a breakpoint in transpiration at a relatively low VPD_leaf_, but a high transpiration rate was kept and a net transpiration increase continued with rising VPD_leaf_ (Fig. 4, Fig. 5). Hence, these genotypes display a more risk taking behaviour, thereby risking hydraulic failure (Sade, Gebremedhin and Moshelion, 2012).

The transpiration reduction at leaf level was significantly correlated to the reduction at whole-plant level, suggesting similar responses to increasing VPD (Fig. 11). The conservative behaviour of genotypes of group I and group II was validated at the whole-plant level by breakpoints in transpiration rate at low VPDs and/or low increases in transpiration afterwards (Fig. 5, Fig. 9).

As demonstrated in other crops, identification of this conservative behaviour towards VPD, opens up possibilities to improve drought tolerance of cultivated banana hybrids. *M. balbisiana* is a parent to many edible bananas belonging to the AAB, ABB and AB genome groups and their subgroups. In line with the conservative behaviour of *M. balbisiana* in response to VPD (Fig 3-4, Fig 6, Fig 8), it has been indicated in many studies that edible bananas with a high portion of B genes are related to drought tolerance (Ekanayake, Ortiz and Vuylsteke, 1994; Thomas, Turner and Eamus, 1998; Turner and Thomas, 1998; Thomas and Turner, 2001; Vanhove *et al*., 2012; Kissel *et al*., 2015; Van Wesemael *et al*., 2018; van Wesemael *et al*., 2019; Eyland *et al*., 2021, 2022; Uwimana *et al*., 2021). Also *M. acuminata spp. zebrina* is a parent to several edible bananas (Carreel *et al*., 2002; Perrier *et al*., 2011; Němečková *et al*., 2018; Baurens *et al*., 2019; Martin, Baurens, *et al*., 2020; Martin, Cardi, *et al*., 2020; Jeensae *et al*., 2021), among others the East-African highland banana subgroup (i.e. Mutika/Lujugira). The East-African highland banana subgroup, endemic to the East-African highlands, is due to its risk taking behaviour sensitive to drought (Kissel *et al*., 2015; van Wesemael *et al*., 2019; Eyland *et al*., 2021; Uwimana *et al*., 2021). Hence, identification of drought tolerance traits in *M. acuminata* ssp. *zebrina* populations provides opportunities to mitigate climate change impacts in this and all other important subgroups. So far, not much is known about the contribution of *M. acuminata* ssp. *errans* to edible bananas. The accession screened in this study and representing *M. acuminata* ssp. *errans*, has been proved to be complex in genome with ancestries coming from ‘*malaccensis*’, ‘*zebrina*’ and ‘*burmannica/siamea*’ (Martin, Cardi, *et al*., 2020).

## 6 Conclusion

The reduction of transpiration response to high VPD is a key trait for water saving and diversity among wild banana relatives was observed. Reductions in transpiration ranging between 37 and 59 %, translated in an increased WUE of 54 to 166 %. *M. acuminata* spp. *errans*, on the one hand, responded most conservative, but was also characterized by low g_*s*_ overall. *M. acuminata* ssp. *zebrina* and *M. balbisiana*, on the other hand, showed strong stomatal closure while maintaining relatively high carbon uptake under low VPD. These two genotypes thus show favourable responses for a specific sub-trait linked to high water use efficiency, providing a potential basis for identification of parent material for drought tolerance breeding.

## Abbreviations

A: photosynthetic rate
A_max_: maximally measured photosynthetic rate
A_meas_: measured photosynthetic rate
ABA: abscisic acid
E_rate_: transpiration rate
E_meas_: measured transpiration rate
E_pred_: predicted transpiration rate
Eq: equation
g_s_: stomatal conductance
h: hour
ITC: International Transit Centre
kPa: kilopascal
L: liter
LA: leaf area
m: meter
min: minutes
mol: moles
m_*tot*_: total weight
PC: principal component
s: seconds
se: standard error
ssp: subspecies
R^2^: R-squared
t_1_: timepoint 1
t_2_: timepoint 2
VPD: vapour pressure deficit
VPD_leaf_: leaf-to-air vapour pressure deficit
_i_WUE: intrinsic water use efficiency
µmol: micromoles
ϕ_E_: transpiration reduction
ϕ_stom_: stomatal reduction

## 7 Supplementary data

**Table S1**: Genotype-specific steady state response of *g*_*s*_, E_rate_, *A* and _i_WUE at increasing VPD

**Table S2**: Genotype-specific transpiration reduction at increasing VPD

**Table S3**: Genotype-specific photosynthetic rate limitation at increasing VPD

**Table S4**: Genotype-specific stomatal reduction

**Table S5**: Genotype-specific whole-plant transpiration rate at increasing VPD

**Table S6**: Genotype-specific whole-plant transpiration reduction

**Fig. S1**: Correlation matrix leaf and whole-plant traits

8 Acknowledgements

The authors would like to thank Edwige Andre for the plant propagation; Hendrik Siongers, Stan Blomme, Loïck Derette and Poi Verwilt for their technical assistance during plant growth and phenotyping.

## 9 Author contributions

SC and RS wrote the concepts for funding. DE performed the experiments and analyzed the data. SC supervised the experiments. SC, CG and DE wrote the manuscript. All authors reviewed and approved the final manuscript.

## 10 Conflict of interest

The authors declare no conflicts of interest.

## 11 Funding

This study was undertaken as part of the initiative ‘Adapting Agriculture to Climate Change: Collecting, Protecting and Preparing Crop Wild Relatives’ which is supported by the Government of Norway. The project is managed by the Global Crop Diversity Trust in partnership with national and international gene banks and plant breeding institutes around the world http://www.cwrdiversity.org/. DE was supported by a scholarship funded by the Global TRUST foundation project ‘Crop Wild Relatives Evaluation of drought tolerance in wild bananas from Papua New Guinea’ [Grant number: GS15024]. CG was supported by a PhD scholarship funded by the Belgian Development Cooperation project ‘‘More fruit for food security: developing climate-smart bananas for the African Great Lakes region”. The authors thank all donors who supported this work also through their contributions to the CGIAR Fund (http://www.cgiar.org/who-we-are/cgiar-fund/fund-donors-2/), and in particular to the CGIAR Research Program Roots, Tubers and Bananas (RTB-CRP) and to the ERA-Net transnational call European Research Projects LEAP Agri H2020 cofund project on food & nutrition security & sustainable agriculture, with funding from national funding agencies for the Project ‘PHENOTYPING THE BANANA BIODIVERSITY TO IDENTIFY CLIMATE SMART VARIETIES WITH OPTIMAL MARKET POTENTIAL IN AFRICA AND EUROPE’.

## 12 Data availability

All data supporting the findings of this study are available within the paper and within its supplementary materials published online

## Notes

### Competing Interest Statement

The authors have declared no competing interest.

## References

Aubert, B. and Catsky, J. (1970) ‘The onset of photosynthetic CO2 influx in banana leaf segments as related to stomatal diffusion resistance at different air humidities’, Photosynthetica, 4(3), pp. 254–256.

Bauer, H. et al.. (2013) ‘The stomatal response to reduced relative humidity requires guard cell-autonomous ABA synthesis’, Current Biology, 23(1), pp. 53–57. doi: 10.1016/j.cub.2012.11.022.

Baurens, F. C. et al.. (2019) ‘Recombination and large structural variations shape interspecific edible bananas genomes’, Molecular Biology and Evolution, 36(1), pp. 97–111. doi: 10.1093/molbev/msy199.

Breshears, D. D. et al.. (2013) ‘The critical amplifying role of increasing atmospheric moisture demand on tree mortality and associated regional die-off’, Frontiers in Plant Science, 4(266). doi: 10.3389/fpls.2013.00266.

Brodribb, T. J. and Jordan, G. J. (2008) ‘Internal coordination between hydraulics and stomatal control in leaves’, Plant, Cell & Environment, 31(11), pp. 1557–1564. doi: 10.1111/j.1365-3040.2008.01865.x.

Carr, M. K. V. (2009) ‘The water relations and irrigation requirements of banana (Musa spp.)’, Experimental Agriculture, 45(3), pp. 333–371. doi: 10.1017/S001447970900787X.

Carreel, F. et al.. (2002) ‘Ascertaining maternal and paternal lineage within Musa by chloroplast and mitochondrial DNA RFLP analyses’, Genome, 45(4), pp. 679–692. doi: 10.1139/g02-033.

Cenci, A. et al.. (2021) ‘Unravelling the complex story of intergenomic recombination in ABB allotriploid bananas’, Annals of Botany, 127(1), pp. 7–20. doi: 10.1093/aob/mcaa032.

Challinor, A. J. and Wheeler, T. R. (2008) ‘Crop yield reduction in the tropics under climate change: Processes and uncertainties’, Agricultural and Forest Meteorology, 148(3), pp. 343–356. doi: 10.1016/j.agrformet.2007.09.015.

Choudhary, S. et al.. (2014) ‘Hydraulic conductance of maize hybrids differing in transpiration response to vapor pressure deficit’, Crop Science, 54(3), pp. 1147–1152. doi: 10.2135/cropsci2013.05.0303.

Christelová, P. et al.. (2017) ‘Molecular and cytological characterization of the global Musa germplasm collection provides insights into the treasure of banana diversity’, Biodiversity and Conservation, 26(4), pp. 801–824. doi: 10.1007/s10531-016-1273-9.

Dai, Z., Edwards, G. E. and Ku, M. S. B. (1992) ‘Control of photosynthesis and stomatal conductance in Ricinus communis L. (castor bean) by leaf to air vapor pressure deficit’, Plant Physiology, 99(4), pp. 1426–1434. doi: 10.1104/pp.99.4.1426.

Devi, M. J., Sinclair, T. R. and Taliercio, E. (2015) ‘Comparisons of the effects of elevated vapor pressure deficit on gene expression in leaves among two fast-wilting and a slow-wilting soybean’, PLoS ONE, 10(10), pp. 1–21. doi: 10.1371/journal.pone.0139134.

Ekanayake, I. J., Ortiz, R. and Vuylsteke, D. R. (1994) ‘Influence of leaf age, soil moisture, VPD and time of day on leaf conductance of various Musa genotypes in a humid forest-moist savanna transition site’, Annals of Botany, 74, pp. 173–178.

Eyland, D. et al.. (2021) ‘The impact of slow stomatal kinetics on photosynthesis and water use efficiency under fluctuating light’, Plant Physiology, 186(2), pp. 998–1012. doi: 10.1093/plphys/kiab114.

Eyland, D. et al.. (2022) ‘High-throughput phenotyping reveals differential transpiration behaviour within the banana wild relatives highlighting diversity in drought tolerance’, Plant Cell and Environment, 45(6), pp. 1647–1663. doi: 10.1111/pce.14310.

Ficklin, D. L. and Novick, K. A. (2017) ‘Historic and projected changes in vapor pressure deficit suggest a continental-scale drying of the United States atmosphere’, Journal of Geophysical Research, 122(4), pp. 2061–2079. doi: 10.1002/2016JD025855.

Fletcher, A. L., Sinclair, T. R. and Allen, L. H. (2007) ‘Transpiration responses to vapor pressure deficit in well watered “slow-wilting” and commercial soybean’, Environmental and Experimental Botany, 61(2), pp. 145–151. doi: 10.1016/j.envexpbot.2007.05.004.

Franks, P. J., Cowan, I. R. and Farquhar, G. D. (1997) ‘The apparent feedforward response of stomata to air vapour pressure deficit: Information revealed by different experimental procedures with two rainforest trees’, Plant, Cell and Environment, 20(1), pp. 142–145. doi: 10.1046/j.1365-3040.1997.d01-14.x.

Franks, P. J. and Farquhar, G. D. (1999) ‘A relationship between humidity response, growth form and photosynthetic operating point in C3 plants’, Plant, Cell and Environment, 22(11), pp. 1337–1349. doi: 10.1046/j.1365-3040.1999.00494.x.

Gaffney, J. et al.. (2015) ‘Industry-scale evaluation of maize hybrids selected for increased yield in drought-stress conditions of the US corn belt’, Crop Science, 55(4), pp. 1608–1618. doi: 10.2135/cropsci2014.09.0654.

Gholipoor, M. et al.. (2010) ‘Genetic variability of transpiration response to vapor pressure deficit among sorghum genotypes’, Field Crops Research, 119, pp. 85–90. doi: 10.1016/j.fcr.2010.06.018.

Grossiord, C. et al.. (2020) ‘Plant responses to rising vapor pressure deficit’, New Phytologist, 226(6), pp. 1550–1566. doi: 10.1111/nph.16485.

Hartigan, J. A. and Wong, M. A. (1979) ‘Algorithm AS 136: A K-Means Clustering Algorithm’, Applied Statistics, 28(1), p. 100. doi: 10.2307/2346830.

Hatfield, J. L. and Prueger, J. H. (2015) ‘Temperature extremes: Effect on plant growth and development’, Weather and Climate Extremes, 10, pp. 4–10. doi: 10.1016/j.wace.2015.08.001.

IPCC et al. (2021) Climate Change 2021: The Physical Science Basis. Contribution of Working Group I to the Sixth Assessment Report of the Intergovernmental Panel on Climate Change. Edited by R. Yu and B. Zhou. Cambridge University Press.

Jeensae, R. et al.. (2021) ‘Cultivar specific gene pool may play an important role in Musa acuminata Colla evolution’, Genetic Resources and Crop Evolution, 68(4), pp. 1589–1601. doi: 10.1007/s10722-020-01088-y.

Jyostna Devi, M., Sinclair, T. R. and Vadez, V. (2010) ‘Genotypic variation in peanut for transpiration response to vapor pressure deficit’, Crop Science, 50(1), pp. 191–196. doi: 10.2135/cropsci2009.04.0220.

Kholová, J. et al.. (2010) ‘Terminal drought-tolerant pearl millet [Pennisetum glaucum (L.) R. Br.] have high leaf ABA and limit transpiration at high vapour pressure deficit’, Journal of Experimental Botany. doi: 10.1093/jxb/erq013.

Kissel, E. et al.. (2015) ‘Transpiration efficiency versus growth: exploring the banana biodiversity for drought tolerance’, Scientia Horticulturae, 185, pp. 175–182. doi: 10.1016/j.scienta.2015.01.035.

Kuromori, T., Sugimoto, E. and Shinozaki, K. (2014) ‘Intertissue signal transfer of abscisic acid from vascular cells to guard cells’, Plant Physiology, 164(4), pp. 1587–1592. doi: 10.1104/pp.114.235556.

De Langhe, E. et al.. (2010) ‘Did backcrossing contribute to the origin of hybrid edible bananas?’, Annals of Botany, 106(6), pp. 849–857. doi: 10.1093/aob/mcq187.

Lawrence, M. G. (2005) ‘The relationship between relative humidity and the dewpoint temperature in moist air: A simple conversion and applications’, Bulletin of the American Meteorological Society, 86(2), pp. 225–234. doi: 10.1175/BAMS-86-2-225.

Lee, Y. H. et al.. (2020) ‘The effect of concurrent elevation in CO2 and temperature on the growth, photosynthesis, and yield of potato crops’, PLoS ONE, 15(10), pp. 1–20. doi: 10.1371/journal.pone.0241081.

Lobell, D. B. et al.. (2013) ‘The critical role of extreme heat for maize production in the United States’, Nature Climate Change, 3(5), pp. 497–501. doi: 10.1038/nclimate1832.

Martin, G., Baurens, F. C., et al. (2020) ‘Chromosome reciprocal translocations have accompanied subspecies evolution in bananas’, Plant Journal, 104(6), pp. 1698–1711. doi: 10.1111/tpj.15031.

Martin, G., Cardi, C., et al. (2020) ‘Genome ancestry mosaics reveal multiple and cryptic contributors to cultivated banana’, Plant Journal, 102(5), pp. 1008–1025. doi: 10.1111/tpj.14683.

Matzner, S. and Comstock, J. (2001) ‘The temperature dependence of shoot hydraulic resistance: Implications for stomatal behaviour and hydraulic limitation’, Plant, Cell and Environment, 24(12), pp. 1299–1307. doi: 10.1046/j.0016-8025.2001.00785.x.

McAdam, S. A. M. and Brodribb, T. J. (2015) ‘The evolution of mechanisms driving the stomatal response to vapor pressure deficit’, Plant Physiology, 167(3), pp. 833–843. doi: 10.1104/pp.114.252940.

McAdam, S. A. M. and Brodribb, T. J. (2016) ‘Linking turgor with ABA biosynthesis: Implications for stomatal responses to vapor pressure deficit across land plants’, Plant Physiology, 171(3), pp. 2008–2016. doi: 10.1104/pp.16.00380.

McAdam, S. A. M., Sussmilch, F. C. and Brodribb, T. J. (2016) ‘Stomatal responses to vapour pressure deficit are regulated by high speed gene expression in angiosperms’, Plant, Cell & Environment, 39(3), pp. 485–491. doi: 10.1111/pce.12633.

Medina, S. et al.. (2019) ‘The plant-transpiration response to vapor pressure deficit (VPD) in durum wheat is associated with differential yield performance and specific expression of genes involved in primary metabolism and water transport’, Frontiers in Plant Science, 9(1994), pp. 1–19. doi: 10.3389/fpls.2018.01994.

Messina, C. D. et al.. (2015) ‘Limited-transpiration trait may increase maize drought tolerance in the US corn belt’, Agronomy Journal, 107(6), pp. 1978–1986. doi: 10.2134/agronj15.0016.

Monteith, J. L. (1995) ‘A reinterpretation of stomatal responses to humidity’, Plant, Cell and Environment, 18(4), pp. 357–364. doi: 10.1111/j.1365-3040.1995.tb00371.x.

Monteith, J. L. and Unsworth, M. H. (2013) ‘Properties of gases and liquids’, in Monteith, J. L. and Unsworth, M. H. (eds) Principles of Environmental Physics. 4th edn. Boston: Elsevier, pp. 5–23. doi: 10.1016/B978-0-12-386910-4.00002-0.

Muggeo, V. M. R. (2008) ‘Segmented: an R package to fit regression models with broken-line relationships’, R news, pp. 20–25.

Němečková, A. et al.. (2018) ‘Molecular and cytogenetic study of East African Highland Banana’, Frontiers in Plant Science, 9(October), pp. 1–13. doi: 10.3389/fpls.2018.01371.

Ocheltree, T. W., Nippert, J. B. and Prasad, P. V. V. (2014) ‘Stomatal responses to changes in vapor pressure deficit reflect tissue-specific differences in hydraulic conductance’, Plant, Cell & Environment, 37, pp. 132–139. doi: 10.1111/pce.12137.

Oren, R. et al.. (1999) ‘Survey and synthesis of intra-and interspecific variation in stomatal sensitivity to vapour pressure deficit’, Plant, Cell & Environment, 22(12), pp. 1515–1526. doi: 10.1046/j.1365-3040.1999.00513.x.

Perrier, X. et al.. (2011) ‘Multidisciplinary perspectives on (Musa spp.) domestication’, Proceedings of the National Academy of Sciences of the United States of America, 108(28), pp. 11311–11318. doi: 10.1073/pnas.1102001108.

Reddy, P. S. et al.. (2017) ‘Molecular cloning and expression analysis of Aquaporin genes in pearl millet [Pennisetum glaucum (L) R. Br.] genotypes contrasting in their transpiration response to high vapour pressure deficits’, Plant Science, 265, pp. 167–176. doi: 10.1016/j.plantsci.2017.10.005.

Richards, R. A. (2000) ‘Selectable traits to increase crop photosynthesis and yield of grain crops’, Journal of Experimental Botany, 51(SPEC. ISS.), pp. 447–458. doi: 10.1093/jexbot/51.suppl_1.447.

Ryan, A. C. et al.. (2016) ‘Gravimetric phenotyping of whole plant transpiration responses to atmospheric vapour pressure deficit identifies genotypic variation in water use efficiency’, Plant Science, 251, pp. 101–109. doi: 10.1016/j.plantsci.2016.05.018.

Sack, L., John, G. P. and Buckley, T. N. (2018) ‘ABA accumulation in dehydrating leaves is associated with decline in cell volume, not turgor pressure’, Plant Physiology, 176(1), pp. 489–493. doi: 10.1104/pp.17.01097.

Sade, N., Gebremedhin, A. and Moshelion, M. (2012) ‘Risk-taking plants: anisohydric behavior as a stress-resistance trait.’, Plant signaling & behavior, 7(7), pp. 767–770. doi: 10.4161/psb.20505.

Sadok, W. and Sinclair, T. R. (2010) ‘Genetic variability of transpiration response of soybean [Glycine max (L.) Merr.] shoots to leaf hydraulic conductance inhibitor AgNO3’, Crop Science, 50, pp. 1423–1430. doi: 10.2135/cropsci2009.10.0575.

Schoppach, R. et al.. (2016) ‘High resolution mapping of traits related to whole-plant transpiration under increasing evaporative demand in wheat’, Journal of Experimental Botany, 67(9), pp. 2847–2860. doi: 10.1093/jxb/erw125.

Sinclair, T. R. et al.. (2010) ‘Assessment across the United States of the benefits of altered soybean drought traits’, Agronomy Journal, 102(2), pp. 475–482. doi: 10.2134/agronj2009.0195.

Sinclair, T. R. et al.. (2017) ‘Limited-transpiration response to high vapor pressure deficit in crop species’, Plant Science, 260, pp. 109–118. doi: 10.1016/j.plantsci.2017.04.007.

Sivasakthi, K. et al.. (2017) ‘Chickpea genotypes contrasting for vigor and canopy conductance also differ in their dependence on different water transport pathways’, Frontiers in Plant Science, 8(1663), pp. 1–16. doi: 10.3389/fpls.2017.01663.

Thomas, D. S. and Turner, D. W. (2001) ‘Banana (Musa sp.) leaf gas exchange and chlorophyll fluorescence in response to soil drought, shading and lamina folding’, Scientia Horticulturae, 90(1–2), pp. 93–108. doi: 10.1016/S0304-4238(00)00260-0.

Thomas, D. S., Turner, D. W. and Eamus, D. (1998) ‘Independent effects of the environment on the leaf gas exchange of three banana (Musa sp.) cultivars of different genomic constitution’, Scientia Horticulturae, 75(1–2), pp. 41–57. doi: 10.1016/S0304-4238(98)00114-9.

Turner, D. W. and Thomas, D. S. (1998) ‘Measurements of plant and soil water status and their association with leaf gas exchange in banana (Musa spp.): a laticiferous plant’, Scientia Horticulturae, 77(3–4), pp. 177–193. doi: 10.1016/S0304-4238(98)00168-X.

Uwimana, B. et al.. (2021) ‘Effect of seasonal drought on the agronomic performance of four banana genotypes (Musa spp.) in the east african highlands’, Agronomy, 11(1). doi: 10.3390/agronomy11010004.

Vadez, V. (2014) ‘Root hydraulics: The forgotten side of roots in drought adaptation’, Field Crops Research, 165, pp. 15–24. doi: 10.1016/j.fcr.2014.03.017.

Vanhove, A.-C. et al.. (2012) ‘Screening the banana biodiversity for drought tolerance: can an in vitro growth model and proteomics be used as a tool to discover tolerant varieties and understand homeostasis’, Frontiers in Plant Science, 3, pp. 1–10. doi: 10.3389/fpls.2012.00176.

van Wesemael, J. et al.. (2019) ‘Using growth and transpiration phenotyping under controlled conditions to select water efficient banana genotypes’, Frontiers in Plant Science, 10, pp. 1–14. doi: 10.3389/fpls.2019.00352.

Van Wesemael, J. et al.. (2018) ‘Homeolog expression analysis in an allotriploid non-model crop via integration of transcriptomics and proteomics’, Scientific Reports, 8(1), pp. 1–11. doi: 10.1038/s41598-018-19684-5.

Williams, A. P. et al.. (2013) ‘Temperature as a potent driver of regional forest drought stress and tree mortality’, Nature Climate Change, 3(3), pp. 292–297. doi: 10.1038/nclimate1693.

Xie, X. et al.. (2006) ‘The identification of genes involved in the stomatal response to reduced atmospheric relative humidity’, Current Biology, 16(9), pp. 882–887. doi: 10.1016/j.cub.2006.03.028.

Yang, Z. et al.. (2012) ‘Temperature effect on transpiration response of maize plants to vapour pressure deficit’, Environmental and Experimental Botany, 78, pp. 157–162. doi: 10.1016/j.envexpbot.2011.12.034.

Zhang, Y. J. et al.. (2013) ‘Midday stomatal conductance is more related to stem rather than leaf water status in subtropical deciduous and evergreen broadleaf trees’, Plant, Cell and Environment, 36(1), pp. 149–158. doi: 10.1111/j.1365-3040.2012.02563.x.

